# Effect of acute physical exercise on motor sequence memory

**DOI:** 10.1101/2020.01.28.922930

**Authors:** Blanca Marin Bosch, Aurélien Bringard, Maria Grazia Logrieco, Estelle Lauer, Nathalie Imobersteg, Aurélien Thomas, Guido Ferretti, Sophie Schwartz, Kinga Igloi

## Abstract

Acute physical exercise improves memory functions by increasing neural plasticity in the hippocampus. In animals, a single session of physical exercise has been shown to boost anandamide (AEA), an endocannabinoid known to promote hippocampal plasticity. Hippocampal neuronal networks encode episodic memory representations, including the temporal organization of elements, and can thus benefit motor sequence learning. While previous work established that acute physical exercise has positive effects on declarative memory linked to hippocampal plasticity mechanisms, its influence on memory for motor sequences, and especially on neural mechanisms underlying possible effects, has been less investigated.

Here we studied the impact of acute physical exercise on motor sequence learning, and its underlying neurophysiological mechanisms in humans, using a cross-over randomized within-subjects design. We measured behavior, fMRI activity, and circulating AEA levels in fifteen healthy participants while they performed a serial reaction time task (SRTT) before and after a short period of exercise (moderate or high intensity) or rest.

We show that exercise enhanced motor sequence memory, significantly for high intensity exercise and tending towards significance for moderate intensity exercise. This enhancement correlated with AEA increase, and dovetailed with local increases in caudate nucleus and hippocampus activity.

These findings demonstrate that acute physical exercise promotes sequence learning, thus attesting the overarching benefit of exercise to hippocampus-related memory functions.

## INTRODUCTION

From typing on our smartphones to tying our shoes, most of our everyday activities engage a wide range of motor skills. Conversely, many neurological disorders such as ataxia (a deficit in motor coordination) or Parkinson’s disease disrupt motor behavior. When learning new motor skills, both explicit and implicit mechanisms are typically involved, and have been shown to rely on hippocampal and striatal circuits ^1, 2^. Moreover, hippocampal damage can thus impair both explicit and implicit memory ^3^. Recently, evidence has accumulated to suggest that regular physical exercise over several weeks or months may promote synaptic plasticity in the hippocampus in animal ^4, 5, 6^ and in human ^7^, while also enhancing learning and memory in general ^8^. Regarding motor skill learning specifically, lower fitness levels have been linked to worse performance on motor sequence learning task ^9^.

Remarkably, one single bout of physical exercise (i.e. acute physical exercise) may also benefit diverse cognitive functions such as executive functions ^10^ but also hippocampal plasticity and function, including declarative memory in humans ^11, 12^. Acute physical exercise increases the levels of endocannabinoids in the brain, which can be measured peripherally ^13, 14^. Higher levels of the endocannabinoid anandamide (AEA) have been positively linked to an increase in cannabinoid receptor 1 (CB1) activity and in hippocampal plasticity ^15, 16^ through long-term potentiation mechanisms ^17^. The hippocampus is not the only brain region modulated by CB1 activity. Recent studies have shown that CB1 activity protects striatal neurons in culture against excitotoxicity ^18^ while CB1 activity is reduced in striatal neurons in a mouse-model of ataxia ^19^. Consistent with enhanced striatal function, one report indicated that acute physical exercise also improved procedural memory in a visuomotor accuracy tracking task in a dose-response manner, with higher exercise intensity significantly more beneficial than lower intensity ^20^. Finally, a recent study reported that overnight consolidation of motor sequence learning was impaired in hippocampal amnesiacs compared to matched controls, despite similar initial learning ^21^, hence further supporting a key contribution of the hippocampus to the consolidation of not only declarative memory but also motor sequences. Based on these observations from different research fields, we hypothesized that one bout of exercise should promote hippocampal (and striatal) activity and functions, plausibly together with increased circulating levels of AEA, and enhance the consolidation of memory for motor sequences. We also predicted that high compared to moderate exercise intensity would yield stronger effects.

To test these hypotheses, we combined behavioral, imaging, and blood samples measurements in an experimental protocol including two conditions of exercise with different intensities (moderate intensity exercise at 70% of maximal heart rate, corresponding to 60% of VO2max and high intensity exercise as 80% of maximal heart rate, corresponding to 75% of VO2max) and one additional no-exercise or rest condition (Fig. 1A). We tested 15 right-handed, healthy and fit males (23.2 ± 4.21 years old) in a randomized within-subjects design involving three visits (one condition per visit). To assess motor sequence learning, we adapted a finger tapping task or serial reaction time task (SRTT) previously used by Ros et al. ^22^, in which participants were trained at executing a sequence of keypresses with the four fingers of their non-dominant left hand as indicated by visual cues on a screen (Fig. 1C). Participants were not told explicitly that they would produce a fixed sequence of 12 button presses (a different sequence was trained on each visit), repeated 10 times in each training block. Thus, at each visit, participants came in the morning and performed the first session (Session 1) of the motor sequence task in the MRI. Session 1 was composed of four blocks (three sequence blocks, and one random block in position 2, Fig. 1B). Then, participants rested for 30 minutes, or cycled at moderate intensity for 30 minutes, or cycled at high intensity for 15 minutes, with an additional 3 minutes of warm-up and 3 minutes of cool-down to limit difference in duration length between exercising conditions and to avoid premature fatigue due to abrupt transition from rest to high intensity exercise. Blood samples were taken right before and right after resting or cycling (Fig. 1A). Forty-five minutes after the second blood sample, i.e. once heart and breathing rates were back to baseline, participants entered the MRI again to perform the second session (Session 2) of the SRTT in the MRI (see Methods section for full description). The effects of exercising on SRTT learning were thus assessed by comparing, for each participant, performance improvement from Session 1 to Session 2 from each exercise condition.

**Figure 1.**
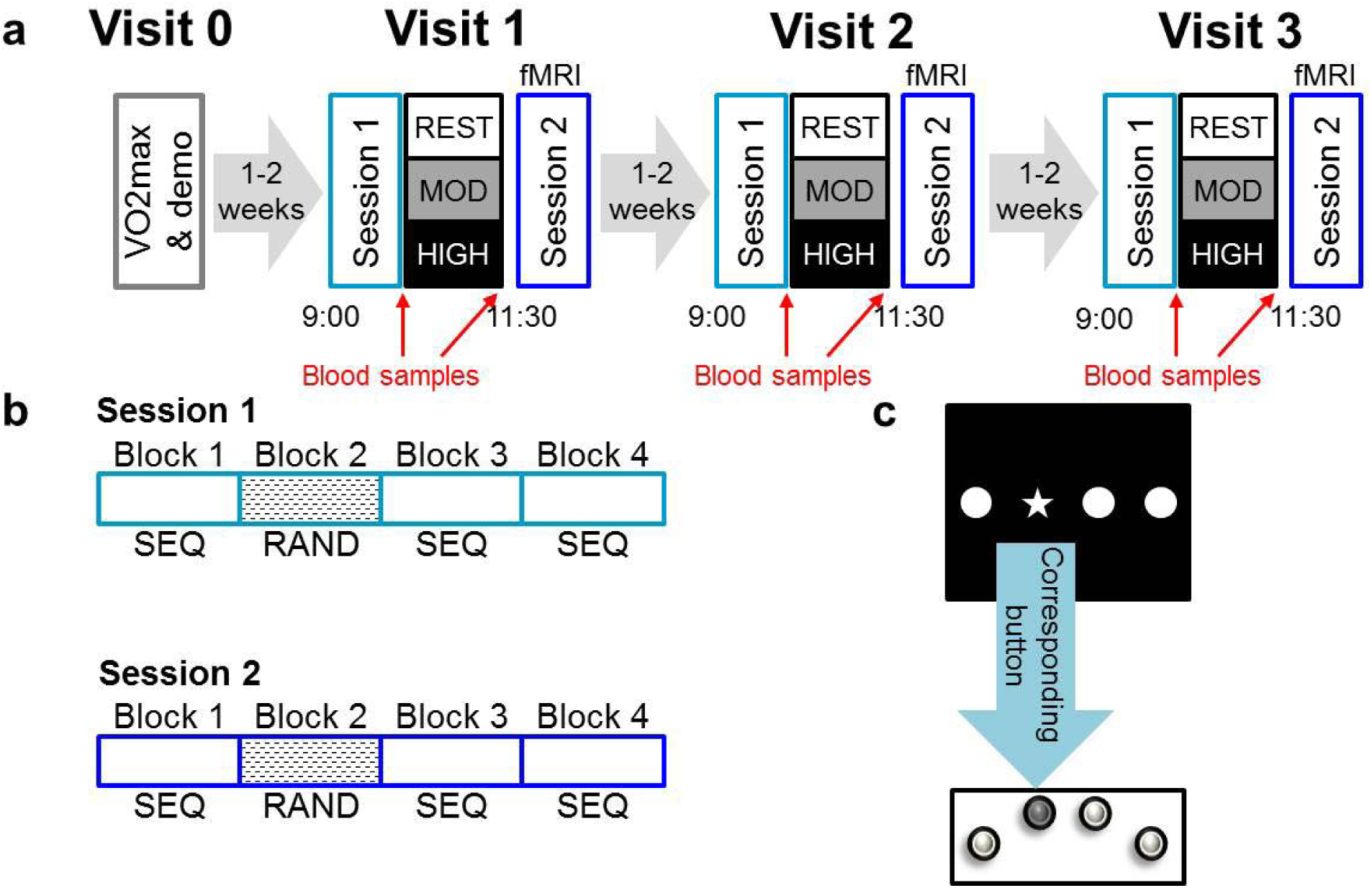
**a)** Experimental protocol. Participants came to the lab four times, first to do a VO2max session and a demo of the SRTT task, then for 3 experimental visits. Each experimental visit consisted of two fMRI sessions (Session1 and Session 2) separated by a Rest, a Moderate exercise or a High exercise session. Right before and right after the Exercise or Rest session blood samples are taken. Experimental visits took part in the morning between 9AM and 12:30PM. **b)** Breakdown of the fMRI sessions, both Session 1 and Session 2 are composed of 4 blocks, i.e. 3 sequence blocks and 1 random block (in position 2) showing 4 blocks for each session. A Block is composed of 120 trials for which participants have to press the button corresponding to the stimuli on the screen. Sequence blocks are composed of 10 repetitions of a 12 element-long sequence of key presses whereas random blocks do not contain any repeated sequence but are matched in keypress frequency and intertrial finger distance to sequence blocks. c) Visual stimulus with corresponding action by participants using the left hand.

## RESULTS

### Behavioral

Participants were instructed to be as quick and as correct as possible on each trial of the SRTT, but there was no time limit. To account for speed-accuracy trade-off, we defined performance as the percentage of correct trials per block divided by the average reaction time on that block. Exercise-related SRTT improvement (or consolidation) was computed as the difference in performance from Session 1 to Session 2, as in Mang et al., 2016^23^. First, we used separate two-sample t-tests against zero and found that participants’ performance improved only for the sequence blocks (rest: T(14)=2.40, p=0.031; moderate: T(14)=3.74, p=0.002; high: T(14)=3.76, p=0.002), but not for the random blocks (rest: T(14)=−0.95, p=0.36; moderate: T(14)=0.39, p=0.70; high: T(14)=1.72, p=0.11). Next, these data were analyzed using a repeated-measures ANOVA with Exercising Condition (rest, moderate and high intensity exercise) and Block type (sequence, random) as within-subjects factors. We report an effect of Exercising Condition (F(2,28)=3.36 p=0.049) and an effect of Block type (F(2,28)=16.28 p=0.001), with no interaction between these effects (F(2,28)=0.43, p=0.655; Fig. 2). To better understand the effect of Exercising Condition, we conducted post-hoc analyses and found a significant difference between rest and high (p=0.049) and a tendency towards significant difference between rest and moderate (p=0.069), while moderate and high did not significantly differ (p=0.558), further there was no significant difference between random blocks according to Exercising condition (p_rest-mod_ = 0.683, p_mod-high_ = 1.000, p_rest-high_ = 0.202). Due to the combination of results, we performed exploratory post-hoc analyses on the non-significant interaction between the effect of Exercising Condition and the effect of Block Type. This analysis revealed no difference between random and sequence blocks in the rest condition (p=0.137), while the difference between random and sequence blocks was significant for both moderate (p=0.009) and high (p=0.007) intensity Exercising conditions.

**Figure 2.**
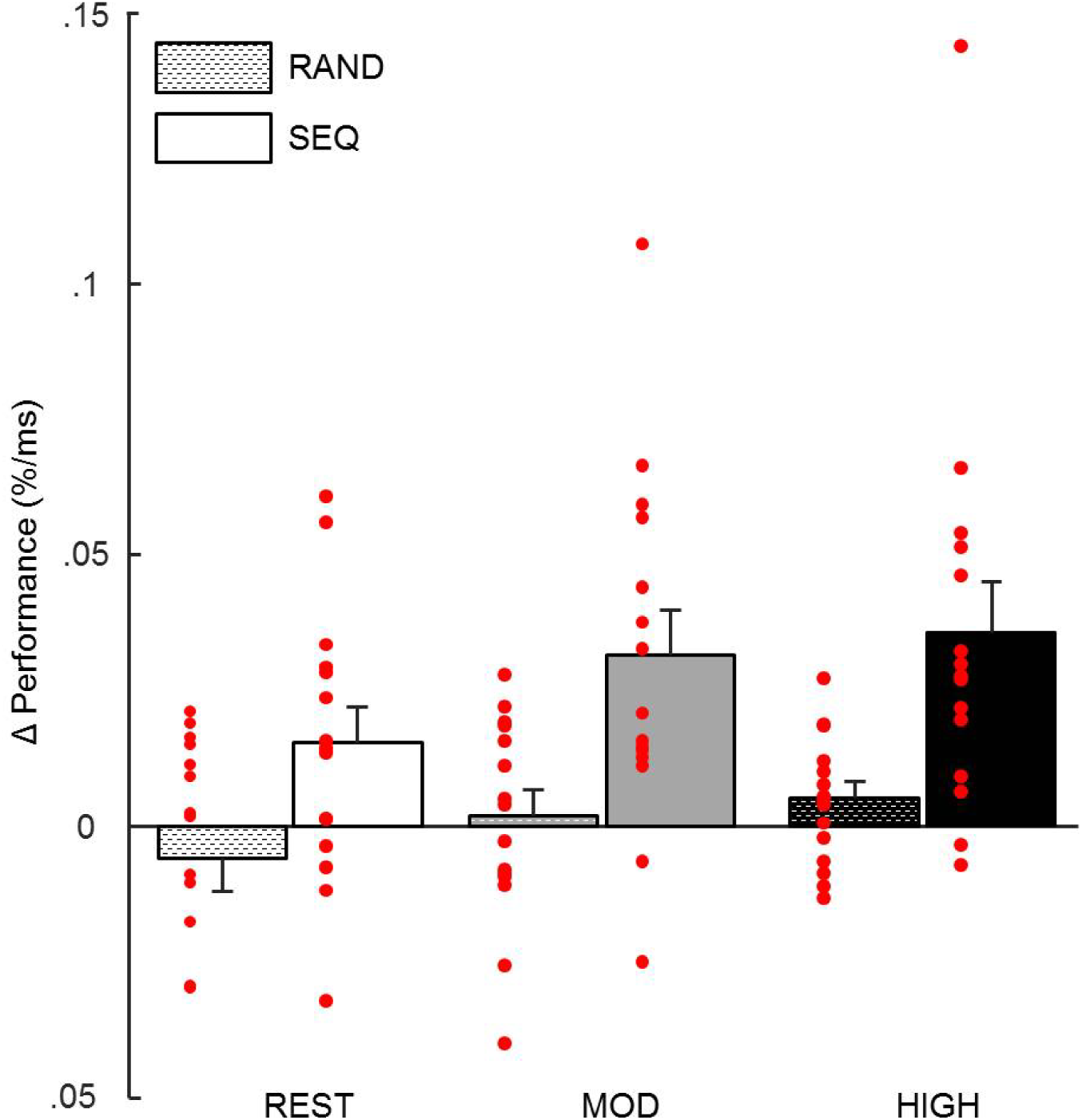
Behavioral results. Performance was defined as mean accuracy (in %) over reaction time (in ms) for both Session 1 and Session 2 of each Exercising condition. Δefficiency is the difference in performance (Δ) from session 1 to session 2. There is a main effect of Exercising condition and of Block type. Post-hoc analyses reveal differences between random and sequence blocks after both moderate and high intensity exercise conditions but not after rest condition. Error bars represent SEM.

This cross-over randomized within-subjects design involved executing a visuo-motor task over three different visits. To assess for possible effects of the order of visit, we computed performance (accuracy/RT) and reaction times for each visit separately and performed two separate repeated-measures ANOVAs for each measurement type, with Visit Order (1-2-3) and Block Type (sequence-random) as repeated measures. For both performance and reaction times, we found no effect of Visit Order (performance: F(2, 28)=0.17, p=0.84; reaction times: F(2, 28)=0.001, p=0.99), an effect of Block Type (performance: F(1, 14)=16.01, p=0.001; reaction times: F(1, 14)=16.79, p=0.001) and no interaction between Visit Order and Block Type (performance: F(2, 28)=1.99, p=0.15; reaction times: F(2, 28)=0.71, p=0.51). To take delay between visits into account, we computed the difference in performance and in reaction times between consecutive visits (difference between visit 2 and visit 1 and difference between visit 3 and visit 2) and added time (in days) between these visits as a cofactor. Here again, these analyses yielded no effect of Visit Order (here difference between consecutive visits), and no interaction with delay between visits (all p>0.50). Altogether, these results suggest that performance or reaction times were not significantly affected by the order of the visits or by the delay between experimental sessions.

### Functional MRI

After standard preprocessing and corrections for breathing and heart rate, the fMRI data from each participant acquired after each exercising condition were analyzed using an event-related general linear model (see Methods section for detailed description). Event-onsets for the sequence and random blocks were defined as the moments when one of the dots on the screen turned into a star. The inverse reaction time (-RT) of each trial was added as parametric modulator of the main event regressors. At the second level, a flexible factorial model was then run with Subject and Block type (sequence and random) as factors, using the contrast images corresponding to the main effects for each Exercising Condition (rest, moderate intensity exercise and high intensity exercise) from each participant. To identify common motor activations between the sequence and random blocks we conducted a conjunction analysis across all participants. This analysis yielded activations in the right precentral gyrus (contralateral to hand used in the task) and left cerebellum, which is consistent with previous literature for this type of tasks ^24^, see Table 1 for all activations). We then tested for global differences in activity elicited after exercise (moderate and high) versus rest for the sequence blocks and found increased bilateral activation of the precuneus (peaks at: [−16 −68 30] and [20 – 60 22]). To directly address one of the main hypotheses that better performance after exercise would rely on hippocampal and striatal circuitry, we looked at the effect of the parametric modulator (-RT) to reveal any regions whose activation increases selectively for faster correct trials, and more so after exercise (moderate and high) than after rest. We found such an effect in the right hippocampus (peak at [30 −6 −24], Fig. 3A) and in the right caudate (peak at [20 24 2], Fig. 3B). Activations in both regions survived small volume correction (cluster level: p_right_ hippocampus=0.002, pright caudate=0.020; peak level: pright hippocampus=0.003, pright caudate=0.038) using predefined regions of interest (see Methods). More importantly, no such activations were found when performing the same analysis on the random blocks. See Table 1 for all activations.

**Table 1.** Activated brain regions at Session 2.

| Main effect during sequence blocks |  |  |  |  |  |  |  |  |  |
| --- | --- | --- | --- | --- | --- | --- | --- | --- | --- |
| Cluster size | Cluster p (unc) | Peak T | Peak Z | Peak p (unc) | x | y | z | Brain Region |  |
| 465 | 2.4E-05 | 9.087 | 5.122 | 1.5E-07 | -26 | -54 | -22 | Left cerebellum exterior |  |
| 507 | 1.3E-05 | 8.851 | 5.062 | 2.1E-07 | -22 | -56 | -50 | Left cerebellum exterior |  |
| 1805 | 4E-12 | 8.122 | 4.864 | 5.7E-07 | 36 | -26 | 46 | Right precentral gyrus |  |
| 422 | 4.7E-05 | 6.402 | 4.308 | 8.3E-06 | -10 | 0 | 54 | Left precentral gyrus |  |
| 182 | 0.003 | 5.426 | 3.918 | 4.5E-05 | 38 | -64 | -4 | Right inf. occipital gyrus |  |
| 70 | 0.049 | 5.080 | 3.763 | 8.4E-05 | 16 | 2 | 50 | Right suppl. motor cortex |  |
| 83 | 0.034 | 5.033 | 3.741 | 9.2E-05 | -34 | 16 | 12 | Left frontal operculum |  |
| Main effect during random blocks |  |  |  |  |  |  |  |  |  |
| 1058 | 6.3E-10 | 8.297 | 4.914 | 4.5E-07 | 42 | -10 | 64 | Right precentral gyrus |  |
| 129 | 0.005 | 7.526 | 4.687 | 1.4E-06 | -32 | 22 | 6 | Left anterior insula |  |
| 356 | 3.6E-05 | 6.754 | 4.434 | 4.6E-06 | -20 | -54 | -46 | Left cerebellum exterior |  |
| 33 | 0.123 | 6.321 | 4.278 | 9.4E-06 | 34 | -42 | -48 | Right cerebellum exterior |  |
| 102 | 0.012 | 6.135 | 4.207 | 1.3E-05 | 34 | 22 | 8 | Right frontal operculum |  |
| 14 | 0.306 | 5.295 | 3.860 | 5.7E-05 | 22 | -64 | 60 | Right sup. parietal lobule |  |
| 11 | 0.365 | 4.875 | 3.667 | 1.2E-04 | -54 | -20 | 30 | Left postcentral gyrus |  |
| 35 | 0.113 | 4.721 | 3.592 | 1.6E-04 | 50 | 6 | 26 | Right precentral gyrus |  |
| 31 | 0.133 | 4.708 | 3.586 | 1.7E-04 | -20 | -8 | 52 | Left precentral gyrus |  |
| 84 | 0.020 | 4.559 | 3.512 | 2.2E-04 | 30 | -40 | 42 | Right sup. parietal lobule |  |
| Conjunction analysis |  |  |  |  |  |  |  |  |  |
| 354 | 4E-04 | 6.540 | 5.053 | 2.2E-07 | -20 | -54 | -50 | Left cerebellum |  |
| 1065 | 6.8E-08 | 6.077 | 4.813 | 7.5E-07 | 38 | -14 | 64 | Right precentral gyrus |  |
| 97 | 0.035 | 5.304 | 4.376 | 6.1E-06 | -32 | 18 | 10 | Bilateral frontal operculum |  |
| 90 | 0.042 | 5.303 | 4.375 | 6.1E-06 | 34 | 20 | 10 |  |  |
| 24 | 0.268 | 4.454 | 3.840 | 6.2E-05 | 32 | -42 | -50 | Bilateral cerebellum exterior |  |
| 118 | 0.022 | 4.439 | 3.830 | 6.4E-05 | -24 | -52 | -22 |  |  |
| 34 | 0.191 | 4.012 | 3.536 | 2E-04 | -8 | 2 | 52 | Left supplementary motor cortex |  |
| Only sequence blocks: Exercise>rest |  |  |  |  |  |  |  |  |  |
| 1263 | 1.5E-12 | 8.629 | 5.004 | 2.8E-07 | -18 | -68 | 24 | Left precuneus |  |
| 159 | 0.001 | 5.228 | 3.831 | 6.4E-05 | -10 | -80 | 10 | Left calcarine cortex |  |
| 7 | 0.427 | 5.077 | 3.762 | 8.4E-05 | 22 | -46 | -52 | Right cerebellum exterior |  |
| 90 | 0.009 | 5.003 | 3.728 | 9.7E-05 | 20 | -60 | 22 | Right precuneus |  |
| Only random blocks: Exercise>rest parametric modulator : inverse reaction time (-RT) |  |  |  |  |  |  |  |  |  |
| 10 | 0.280 | 5.39 | 3.90 | 4.7E-05 | -42 | -86 | -4 | Left inferior occipital gyrus |  |
| Only sequence blocks: Exercise>rest parametric modulator : inverse reaction time (-RT) |  |  |  |  |  |  |  |  |  |
| Cluster size | Cluster p(unc) | Peak T | Peak Z | Peak p (unc) | SVC p-val | x | y | z | Brain Region |
| 88 | 0.006 | 6.822 | 4.457 | 4.15E-06 | 0.002 | 30 | -6 | -24 | <u>Right hippocampus</u> |
| 27 | 0.099 | 5.219 | 3.826 | 6.5E-05 | 0.02 | 22 | 24 | 6 | <u>Right caudate nucleus</u> |

**Table 2.** Individual physical exercise parameters.

| Subject | Antropometric data |  |  | Maximal data |  |  |  | Secondary criteria |  |  |  |  |  | Secondary ventilatory threshold VT2 (W) | High intensity exercise |  | Moderate intensity exercise |  |
| --- | --- | --- | --- | --- | --- | --- | --- | --- | --- | --- | --- | --- | --- | --- | --- | --- | --- | --- |
|  | age | Height (cm) | Weight (kg) | VO2max (L/min) | VO2max (ml/kg/min) | Power (W) | Maximal heart rate (bpm) | VO2 plateau | Respiratory Exchange Ratio | 220-age | 210-(0.65 x age) | 202-(0.72 x age) | 207-(0.68 x age) |  | Estimated P_75% VO2max (W) | Actual high intensity | Estimated P_60% VO2max (W) | Actual moderate intensity |
| 1 | 22.7 | 182.5 | 76.5 | 4.8 | 62.7 | 400 | 184 | YES | 1.12 | 197 | 195 | 187 | 192 | 375 | 250 | 250 | 200 | 180 |
| 2 | 20.9 | 176 | 79.5 | 3.7 | 46.5 | 250 | 201 | YES | 1.05 | 199 | 196 | 188 | 193 | 225 | 175 | 175 | 125 | 110 |
| 3 | 33.2 | 176 | 62 | 2.5 | 40.3 | 225 | 183 | NO | 1.11 | 187 | 188 | 179 | 184 | 200 | 150 | 139 | 125 | 108 |
| 4 | 22.5 | 180.5 | 68.2 | 4.1 | 60.1 | 375 | 181 | YES | 1.01 | 198 | 195 | 187 | 192 | 325 | 225 | 225 | 160 | 173 |
| 5 | 20.7 | 188 | 87.2 | 3.5 | 40.1 | 300 | 196 | YES | 1.10 | 199 | 197 | 188 | 193 | 275 | 175 | 154 | 140 | 140 |
| 6 | 21.9 | 183.5 | 75.5 | 3 | 39.7 | 325 | 198 | YES | 1.09 | 198 | 196 | 187 | 192 | 250 | 200 | 204.5 | 150 | 157 |
| 7 | 21.6 | 180.5 | 78.5 | 3.8 | 48.4 | 325 | 191 | YES | 1.10 | 198 | 196 | 187 | 192 | 300 | 225 | 225 | 175 | 189 |
| 8 | 34.6 | 175.5 | 73.5 | 3.9 | 53.1 | 350 | 182 | YES | 1.11 | 185 | 187 | 178 | 183 | 300 | 210 | 221 | 160 | 178.3 |
| 9 | 24.9 | 187.5 | 84.5 | 4.2 | 49.7 | 350 | 185 | YES | 0.99 | 195 | 194 | 185 | 190 | 275 | 225 | 228 | 175 | 175 |
| 10 | 23.1 | 179.5 | 70 | 3 | 42.1 | 275 | 195 | YES | 1.18 | 197 | 195 | 186 | 191 | 275 | 155 | 162 | 130 | 130 |
| 11 | 23.1 | 177 | 69.5 | 4 | 57.6 | 325 | 186 | YES | 1.04 | 197 | 195 | 186 | 191 | 275 | 225 | 225 | 175 | 175 |
| 12 | 23.1 | 179.5 | 68 | 3.4 | 50.0 | 325 | 185 | YES | 1.13 | 197 | 195 | 186 | 191 | 300 | 200 | 214 | 150 | 157 |
| 13 | 22.9 | 178.5 | 67.2 | 3.3 | 49.1 | 300 | 199 | YES | 1.14 | 197 | 195 | 186 | 191 | 275 | 175 | 164 | 140 | 140 |
| 14 | 21.1 | 177.5 | 75.5 | 4.1 | 53.6 | 330 | 202 | YES | 1.02 | 199 | 196 | 188 | 193 | 250 | 200 | 188 | 150 | 137.3 |
| 15 | 20.4 | 176.5 | 67 | 3.2 | 47.0 | 315 | 190 | YES | 1.06 | 200 | 197 | 188 | 193 | 250 | 200 | 170 | 140 | 134 |
| MEAN | 23.8 | 179.9 | 73.5 | 3.6 | 49.3 | 318.0 | 190.5 |  | 1.08 | 196.2 | 194.5 | 185.8 | 190.8 | 276.7 | 199.3 | 196.3 | 153.0 | 152.2 |
| SD | 4.3 | 4.0 | 7.0 | 0.6 | 7.2 | 44.8 | 7.4 |  | 0.05 | 4.3 | 2.8 | 2.9 | 2.9 | 41.7 | 28.7 | 33.7 | 21.4 | 25.9 |

**Figure 3.**
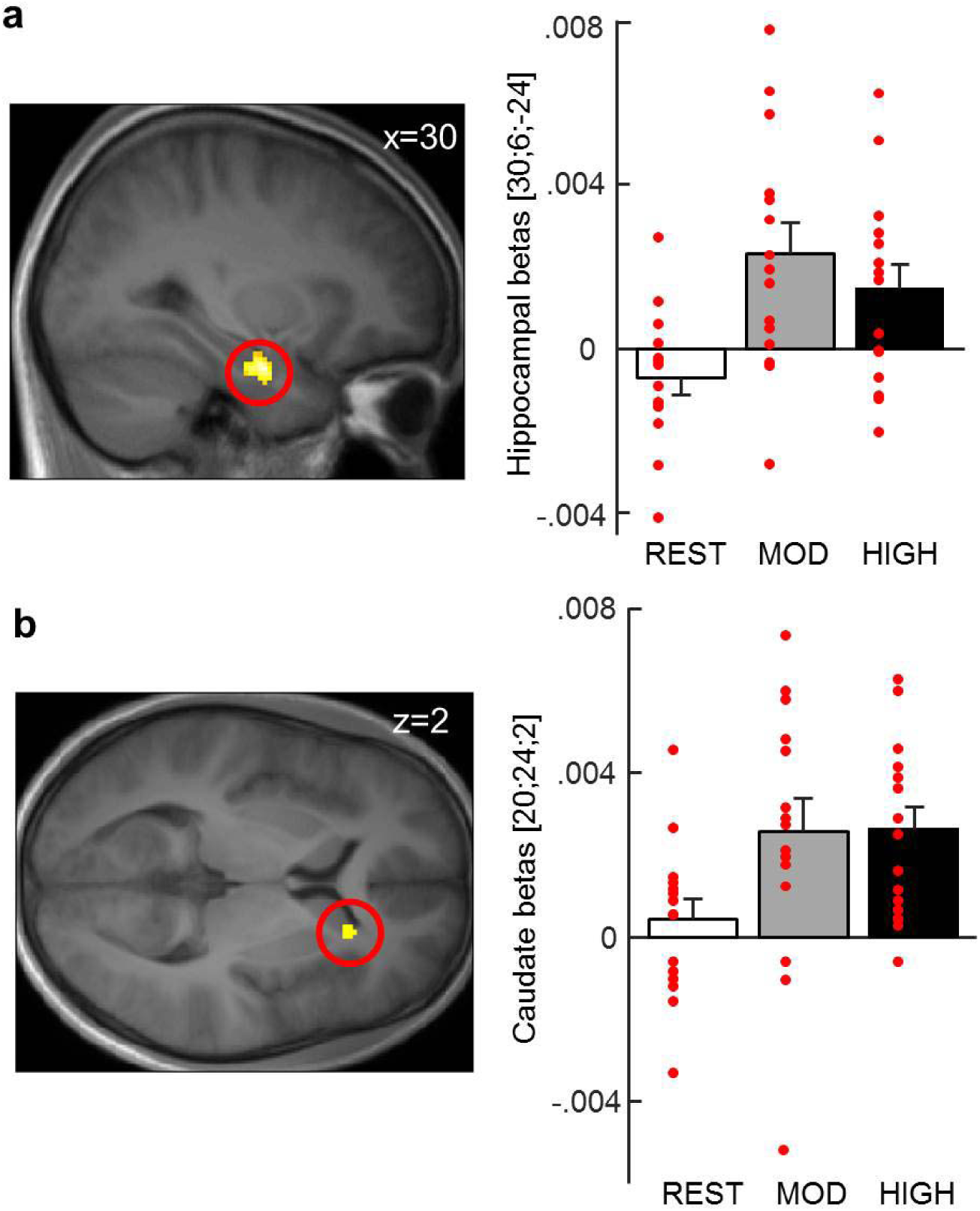
Increased activity in right hippocampus and right caudate for EXERCISE > REST pmod –RT during Session 2. Left: fMRI activations in **a)** right hippocampus (peak at [30; −6; −24]) and **b)** right caudate nucleus (peak at [20; 24; 2]) during Session 2 of the SRTT task after moderate and high intensity Exercising Conditions versus the rest Exercising Condition, with reaction times as trialwise parametric modulator. Right: betas for the fMRI activations. All reported activations survive SVC correction for a small volume defined from the AAL atlas in the WFU PickAtlas toolbox (Wake Forest University School of Medicine) for SPM12. For display purposes, activations are thresholded at p<0.005. Error bars represent SEM.

### Anandamide

Blood samples were taken before (baseline) and after each exercising condition. Analyses on the difference between the second and the first baseline sample revealed a significant effect of Exercising Condition on AEA levels (F(2, 28)=41.991, p<0.001), with increased AEA after moderate and high intensity exercise compared to after rest (p_mod-rest_<0.001; p_high-rest_<0.001) and no difference between both exercising conditions (p_high-mod_=0.080; Fig. 4A).

**Figure 4.**
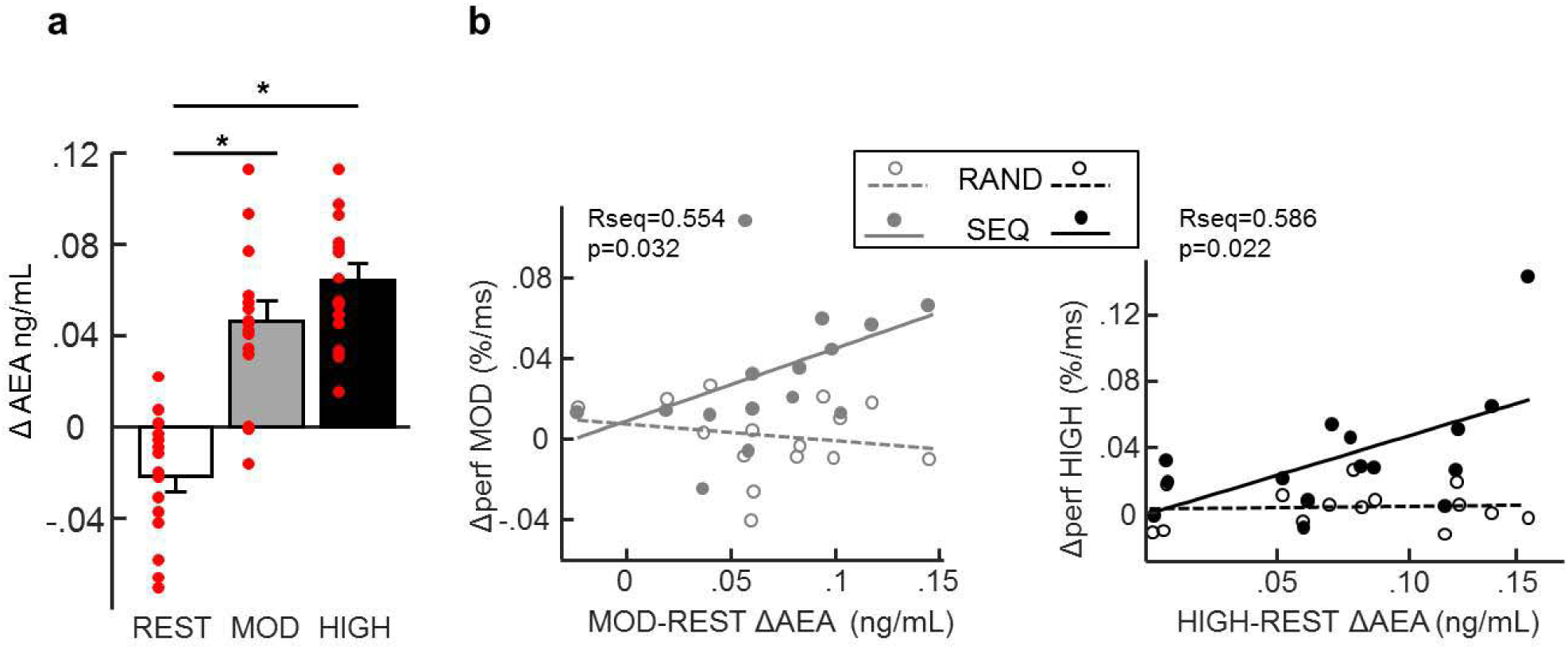
**a)** Increased Anandamide level (AEA) after moderate and high physical exercise compared to after rest. For all Exercising Conditions Δ AEA corresponds to the difference in AEA between the second blood sample taken after exercise or rest and the first blood sample taken before exercise or rest. Error bars represent SEM. **b)** Left panel: Correlation between the increase in performance from Session 1 to Session 2 (Δ performance) for the moderate intensity exercising condition and the increase in Anandamide levels from rest to moderate Exercising conditions (MOD-REST ΔAEA) for the sequence and the random Blocks. Correlation is significant in sequence Blocks but not in the random Blocks. Right panel: Correlation between the increase in performance from Session 1 to Session 2 (Δ performance) for the high intensity exercising condition and the increase in Anandamide levels from rest to high Exercising conditions (HIGH-REST ΔAEA) for the sequence and the random Blocks. Correlation is significant in the sequence Blocks but not in the random Blocks.

Additionally, we found positive correlations between AEA increase and performance on the sequence blocks for both the moderate (R=0.554 p=0.032) and for the high (R=0.586 p=0.022) intensity exercising conditions (Fig. 4b). AEA changes did not correlate with performance on the random blocks (R= −0.23 p=0.41 for moderate condition and R= 0.07 p=0.81 for the high condition). We also directly compared the correlation coefficients between sequence and random blocks for moderate and high conditions using Fisher’s r to z method. We obtained a significantly different correlation during the moderate intensity condition (Fisher z=2.1, p one-tailed =0.018, p two tailed =0.036) and a trend for the high intensity condition (Fisher z=1.48 p one tailed=0.069, p two tailed = 0.139). These results suggest a link between performance for the sequence blocks and increase in AEA.

Finally, as exploratory analyses, we tested whether changes in AEA levels correlated with hippocampal or caudal activations, but did not observe any such correlation (all p> 0.10).

## DISCUSSION

This study shows that one short bout of acute physical exercise at both moderate and high intensities increases AEA levels and hippocampal and striatal recruitment during SRTT, dovetailed with benefits for motor learning performance.

At the behavioral level, participants successfully learned the sequence, as shown by their better performance for sequence compared to random blocks. We also observed a significant overall effect of acute physical exercise, indicating a clear benefit from engaging in high intensity exercise and tending towards an effect for moderate intensity exercise rather than resting. Because we observed a main effect of Exercising condition and no interaction between Exercising Conditions and Block type, this suggests that exercise may have a general positive impact on visuo-motor discrimination processes. However, the participants’ performance on the random blocks did not improve in any of the Exercising Conditions. This pattern of results alludes to some differential impact of exercise on memory processes, beyond mere visuo-motor adaptation effects. We thus conducted a post-hoc exploration within each exercising condition, in an attempt to better disentangle the effects of exercise on motor skill learning from those affecting general visuo-motor processes. This analysis showed significant performance improvement for the sequence (compared to the random) blocks only after acute exercise (moderate and high) but a non-significant trend after the rest condition. Because the results from the brain imaging analyses revealed a strong link between SRTT performance and hippocampal (and striatal) activity after exercise (see also discussion below), we consider that the trend observed in the behavior as providing further converging support to the hypothesis that exercise promotes the consolidation of motor sequence memory.

Results from the literature present some inconsistencies, with acute exercise improving motor skill learning in two within-subjects studies ^23, 25^, and in a dose-dependent manner in a between-subjects study ^20^, but also no effect in yet another between-subjects study ^26^ and one non-significant increase in a further between-subjects study^27^. Please note that between-subjects designs may not have provided sufficient sensitivity for this type of manipulation. In particular, in our within-subjects study, we used two very carefully measured and individually-calibrated exercise intensities. While high intensity exercise yielded numerically better performance, we did not find evidence for any dose-dependent effect, with an overall exercising effect but no significant difference between moderate and high intensity exercise. Most studies showing an effect of acute physical exercise on motor skill learning only compared one intensity (which is usually high) to rest ^23, 26, 27, 28, 29^. The previously mentioned study by Thomas et al. ^20^ is one of the few comparing more than one exercise intensity to rest. However, this is an interval-based exercising intervention with alternating 2-3 minute blocks of exercise at a given intensity (EX45 condition – more or less corresponding to our moderate condition or EX90 condition – which is very high intensity a measure they consider comparable to an effort at maximal heart rate) We fully agree with these authors’ statement that exercise intensity plays a crucial role; here we complement their approach with different exercising intensities at one continuous Exercising condition, which is more suitable for rehabilitation trainings and more easily implementable for patient groups.

At the brain level, we found increased bilateral precuneus activity for both the moderate and high intensity exercising conditions as compared to the rest condition, consistent with the role of the precuneus in motor tasks requiring coordination and/or imagery ^30, 31^. Next, we report that activity in the right hippocampus and caudate nucleus covaried with behavioral performance on the SRTT (faster correct trials) after both moderate and high intensity exercise (compared to rest), thus directly confirming our main initial hypothesis that hippocampal and striatal circuits may be involved in exercise-related performance gains. No such relationship was found for the random sequences, further suggesting that these regions contributed to motor sequence learning, beyond simple visuo-motor mapping,

These results are consistent with motor sequence learning incorporating both implicit and explicit learning components, which implicate striatal and hippocampal regions ^32, 33^. Moreover, a positive relationship between activity in the striatum and hippocampus and motor skill learning was also observed in Parkinson’s patients and healthy controls after a 3-month long exercising protocol ^34^. While the results of our study suggest that long-term regular exercise and one single session of exercise may impact the same brain regions, the exact mechanisms underlying the observed brain-behavior relationships are likely to differ ^35^. Specifically, in our case, activity in the hippocampus and striatum appeared to contribute to response times for accurate motor output, while after long-term training in ^34^ this effect was seen in Parkinson’s patients, but not in controls because of a ceiling effect.

Here we additionally tested whether changes in SRTT performance may relate to exercise-related changes in endocannabinoids’ levels (i.e. AEA). First, we replicated a significant AEA increase in response to physical exercise ^13, 14^. Note that levels slightly dropped after rest, as expected from metabolic and circadian fluctuations ^36, 37^. Second, we found a positive correlation between AEA levels and SRTT performance for both the moderate and for the high intensity exercising conditions, suggesting that sequence learning may have benefited from exercise possibly mediated at least in part by AEA increase. Please note that while both AEA increase and higher hippocampal/striatal activity were linked to increased SRTT performance (for the sequence Block in the moderate and high Exercising Conditions), both measures did not correlate (AEA correlation with hippocampal activity: p_rest_=0.101, p_mod_=0.308, p_high_=0.168; AEA correlation with caudal activity: p_rest_=0.354, p_mod_=0.260, p_high_=0.970). Because of such joined relationship with performance and because the existing animal literature strongly supports that AEA promotes plasticity in both the hippocampus and the striatum ^17, 38, 39^, we cannot rule out that increased AEA affected hippocampal and/or striatal function in our experiment, despite the absence of a simple linear relationship. Moreover, AEA is the strongest endogenous agonist of the CB1 system ^40^, which itself has known neuroprotective effects ^41^.

### Limitations

There are several limitations in this study, among which we here list a few. First, although we estimated our sample size from another published study^22^ using the exact same task, we acknowledge that our estimation may be optimistic (i.e. relatively small), this is why we used a within-subjects design to compensate for possible lack of statistical power. Second, based on previous literature, moderate intensity exercise was set at 30 minutes whereas high intensity lasted 21 minutes (including warm-up and cool-down) to avoid exhaustion. Thus, the durations of the moderate and high intensity exercise conditions slightly differed to achieve comparable overall effort demands and exhaustion levels. Third, concerning the behavioral results: while we observed a significant main effect of Exercising Condition, the comparison between high intensity exercise and rest was significant, but the comparison between moderate intensity exercise and rest yielded a non-significant trend (p=0.069). Moreover, although all performance measures were defined as the difference between performance in Session 2 as compared to Session 1, and Exercising conditions carefully randomized, repeating a motor task may be associated with some unspecific learning across subsequent visits. However, we did not find any effect of visit order neither for performance nor for reaction times. Fourth, the delay in days between experimental visits could not be maintained constant because of external factors (availability of the scanner, of the nurse, and of the physical exercise expert together with the availability of the participant), but the delay did not affect performance or reaction times either. Please note that the time between Session 1 and Session 2 was always constant and all experimental visits were performed in the morning respecting a constant circadian timing. Lastly, here we investigated AEA levels in relation to motor sequence learning, because AEA is reportedly involved in hippocampal plasticity mechanisms and modulated by exercise. However, although we observed a promising positive correlation between motor sequence learning and concentration changes in this molecule, there are likely many other molecules involved in the effect of exercise on motor sequence learning.

To conclude, here we demonstrate that exercise enhanced hippocampal and striatal function and increased endogenous endocannabinoids levels, and that both were linked to better behavioral performance. The neurophysiological mechanisms highlighted in this study provide additional evidence that physical exercise could possibly prevent cognitive decline, especially when hippocampal function is affected. Although more research is still needed, another plausible implication of the present findings concerns people who are unable to perform physical activity on their own due to disease, disability or old age, and who could benefit from pharmacotherapies targeting the AEA or CB1 system as a potential alternative to exercise.

## METHODS

### Participants

Twenty-one healthy young males between the ages of 18 and 35 participated in this study, for which they gave informed consent and were financially compensated. This study was approved by the Ethics Committee of the Geneva University Hospitals. All methods were carried out in accordance with relevant guidelines and regulations. All participants included in this study were right-handed, non-smokers, without psychiatric or neurological history, and had a normal or corrected-to-normal vision. None of the participants had any musical training, which may imply the learning of finger motor sequences, potentially similar to the SRTT. Participants scored within normal ranges on self-assessment questionnaires for depression BDI, ^42^, anxiety STAI, ^43^, and circadian typology ^44^.

All the included participants had a VO2max above 40ml/kg/min and below 65ml/kg/min, which represent values observed in recreationally active individuals, see VO2max measure for details below.

This yielded a homogenous sample of regularly exercising young men. One participant had to be excluded for non-compliance and five were excluded due to technical difficulties. The resulting 15 participants had a mean age of 23.699 ± 4.024.

Concerning the sample size, please note that although 15 participants may appear at the lower limits, here we used a carefully controlled cross-over within-subjects design, which is by essence less affected by inter-individual variability than between-subjects studies. Between-subjects studies typically require 4 to 8 times more participants than within-subjects designs ^45^, and even with half the number of participants within-subjects studies are much more powerful than their between-subjects counterpart ^46^. Based on the effect sizes from a previous behavioral study using the same SRTT as administered here, we determined that the minimum sample size of a within-subjects design for a targeted power of over 0.95, was 6 ^22^. Concerning the fMRI analyses, based on data from previous studies using finger sequence learning ^32, 33^, we established that the minimum sample size for activation in the hippocampus and/or striatum was 7.

### Experimental procedure

Participants came to the lab four times, for one introductory visit (visit 0) and three experimental visits (visits 1, 2 and 3).

### Visit 0

Participants performed a VO2max procedure (detailed below), as well as a demo session of the serial reaction time task (SRTT). Only those participants with a VO2max between 40 and 65 ml/kg/min were called back and invited to complete the rest of the study. These VO2max limits were set specifically so that only physically trained participants were included, thus excluding sedentary participants or elite athletes.

### Visits 1, 2 and 3

These visits were spaced by about two weeks, and exercising conditions and task versions (see below) were fully counterbalanced, according to a cross-over within-subjects design.

Participants were instructed to keep a regular exercising schedule during the week before each visit, and their exercising schedule was documented using a fitness tracker (Fitbit Charge HR, Fitbit, San Francisco, USA). They were also instructed to avoid intense physical activity during the 48 hours preceding each experimental visit.

For each visit, the same schedule was followed, starting at 8:00 AM and ending at 12:00 PM (Figure 1A). Participants first had a standardized breakfast in the lab including black coffee or tea, orange or apple juice, bread and jam. They were allowed to have as much of these items as they wanted, as soon as they would have the same amount on all visits. This breakfast minimized fat intake, because the latter is known to potentially affect circulating levels of the endocannabinoid anandamide (AEA). One hour and a half later, participants were placed in the MRI scanner and performed the first session of the SRTT, after which a medical doctor took a first blood sample. Participants were then asked to rest or exercise while wearing a Polar RS800CX N (Polar RS 800 CX, Polar, Finland) to measure heart rate. In the case of exercise, participants pedaled on a cycle ergometer (Ergoline GmbH, Bitz, Germany), maintaining a pedaling frequency between 60 and 80 cycles per minute. Exercise intensities were determined individually by adapting the load on the ergometer so as to match 70% (moderate intensity) or 80% (high intensity) of each participant’s maximum heart rate. For moderate intensity participants pedaled for 30 minutes while for high intensity exercise we reduced this time to 15 minutes, but included a 3 minutes warm-up and 3 minutes cool-down period amounting to a total of 21 minutes of cycling. This was done to avoid exhaustion as here our main goal was to look at the effect of high physical exercise, and maintaining an effort at 80% of maximal heart rate for 30 minutes is highly demanding and may not be achieved by all participants. Even more important for the present study, 30 mins of exercise at 80% of maximal heart rate would have significantly increased stress level in all participants as reviewed in ^47^, which in itself may compromise memory consolidation ^48, 49^. Therefore, we limited the duration of the high intensity session. On the other hand, based on previous research, including ours, we knew that effects for the moderate intensity exercise may require some sustained exercise session ^11, 20^. Hence, to achieve two exercise sessions of different intensities, but comparable overall effort demands and exhaustion levels, we had to slightly adapt the session durations.

For the ‘rest’ condition, participants sat on a chair and were allowed to look through selected magazines. Shortly after the end of the exercising or rest period, a second blood sample was taken. To allow for breathing and heart rates to go back to baseline, participants waited for 45 minutes before they were placed again inside the MRI for the second session of the SRTT.

### Serial Reaction Time Task (SRTT)

We used the SRTT task described in Ros et al. ^22^. In each experimental visit (i.e. Visits 1, 2, and 3), participants performed two sessions of the SRTT, one before and one after the exercising or rest period, and each composed of 4 blocks (Fig. 1B). Participants were shown a black screen with four white horizontally-organized dots corresponding to their index, middle, ring and little fingers of their non-dominant hand (Fig. 1C). Each finger was placed over a button of a 4-button MRI-compatible box (HH-1×4-CR, Current Designs Inc.). When one of the dots turned into a star, the participants had to press the corresponding button as fast as they could. As soon as they pressed a button, another dot would turn into a star and they had to then press the button associated to the new star, and so on, until the full SRTT session was completed.

In each sequence block, unbeknownst to them, the button presses formed a sequence of 12 presses (or trials), repeated 10 times. Random blocks were designed to match sequence blocks with respect of the number of digit presses and digit span between trials, but there was no underlying repeated sequence. Each sequence or random block was formed of 120 trials, and each session comprised 4 blocks, with the random block at the second position. None of the participants spontaneously reported to have noticed any repetition during the sequence blocks and they were not debriefed afterward either. One distinct 12-item sequence (or version of the task) was used in each visit (randomly assigned to exercising or rest conditions), and we did not observe any significant effect of task version on SRTT performance (F(2,28)=0.690 p=0.510). Consolidation time between Session 1 and Session 2 of the SRTT always amounted to 2 hours and 30 minutes, please note that differences in time to complete the task were very small (mean: 34.5+/−5.3 sec), and were very little comparatively to the duration of the consolidation time and should therefore not have influenced this measure.

During Visit 0 participants did a small introductory session of the task (demo version), where all trials were random and feedback was given after every trial (correct or not).

### Behavioral analysis

The program Statistica (Version 12, www.statsoft.com, StatSoft, Inc. TULSA, OK, USA) was used for all behavioral analyses performed in this study. Performance was defined as accuracy (percentage of correct responses) divided by reaction time (mean reaction time per correct trial), to accounts for speed-accuracy trade-off effects, as typically observed in speeded reaction time tasks. To remove any potential initial performance difference for a given sequence, we compared pre and post exercising condition (Session 1 to Session 2) for both sequence and random blocks to be able to separately assess sequence-related learning effects and general visuo-motor learning effects (as in Mang et al. 2016^23^ and Meehan et al. 2011^50^), therefore we ran our analyses on the difference in performance from Session 1 to Session 2. To determine whether learning occurred from Session 1 to Session 2, we performed t-tests against zero for each relevant variable. Then, we performed a repeated-measures ANOVA to test for effects of exercise on motor sequence learning using an ANOVA with Exercising Condition (rest, moderate intensity, high intensity) and Block type (random, sequence) as factors. All post-hoc analyses were performed using the Bonferroni post-hoc method and all correlations were Spearman Rank Order correlations.

### VO2max measure

During Visit 0, all participants performed a maximal incremental test on an electrically braked cycle ergometer (Ergometrics er800S, Ergoline, Jaeger, Germany). Using a metabolic unit (K4b^2^, Cosmed, Italy), respiratory gas flow and ventilation were measured breath-to-breath. This metabolic unit was composed of a Zirconium Oxygen analyzer, an infrared CO2 meter and a turbine flowmeter. The gas analyzers and the turbine were calibrated as recommended by the manufacturer, the former with ambient air and with a mixture of known gases (O_2_ 16%, CO_2_ 5%, N_2_ as balance), and the latter using a 3-L syringe. Heart rate was monitored on a beat-by-beat basis using cardiotachography (Polar RS 800 CX, Polar, Finland).

The following gas exchange variables were recorded on a breath-by-breath basis: V^▪^O2, V^▪^CO2, V^▪^E, and respiratory exchange ratio. They were then averaged over 10–second sliding intervals for later analysis. The test started with participants pedaling at a power output of 50W for 4 minutes. Power was increased by 25W every 2 minutes until achieving 80% of maximal heart rate predicted by age. From this point forward, power was increased by 25W every 1 minute until volitional exhaustion. Pedaling frequency had to be maintained in the range of 60 – 80 cycles per minute throughout the test.

We used the criteria recommended by the American Thoracic Society (ATS) /American College Chest Physicians (ACCP) statement on cardiopulmonary exercise testing^51^ to ascertain VO2max. VO2max was determined when the following criteria were fulfilled: respiratory exchange ratio > 1.1, plateau in VO2 (change of <100 mL·min^−1^ in the last three consecutive 20-s averages), and a heart rate less than 10 beats·min-1 away from the calculated maximal level predicted by age. The results from this test were used to select the appropriate power output for the experimental visits, based on the relationship between power output and VO2.

From our maximal incremental effort test, we first determined individual VO2max and maximal heart rate levels. These parameters were then used to define exercising intensities during moderate and high intensity Exercising conditions: we set the initial work rate (i.e. power output developed by the participant in Watts, against the resistance imposed by the cycle ergometer) based on individual VO2max and adjusted this work rate based on heart rate levels (see the Parameters of physical exercise during Exercising conditions for details). Additionally, to insure that all participants remained in the same intensity domain, data of the incremental effort tests were analysed for determination of the second ventilatory threshold (VT2). VT2 was determined by the ventilatory equivalent method, where a clear breakpoint occurs on the plot between the increase in ventilation to the increase in CO2 and partial pressure of exhaled carbon dioxide decrease ^52, 53^.

We only included participants whose VO2max was above 40ml/kg/min and below 65ml/kg/min, which correspond to values observed in recreationally active individuals. The lower bound was used to ensure that participants would tolerate the high intensity condition, and specially would be able to continue exercising despite the uncomfortably high ventilation rate associated with exercise at intensities close to the second ventilatory threshold (VT2). The upper bound was needed for a homogeneous high intensity exercise condition that remains below VT2 (which corresponds in a cycling paradigm to about 75-80% of VO2max or 85-90% of maximal cardiac frequency for moderately fit participants). In very highly endurance trained individuals, whose VO2max is above 65ml/kg/min VT2 is often higher, at about 80% of VO2max or even higher in to class athletes, corresponding to 90% (or even higher) of maximal cardiac frequency. Thus, an intensity defined as 75% of VO2max would not correspond to comparable difficulty levels for moderately fit vs. highly trained participants, whether it is below or above VT2. In all of the 15 participants, both work rate corresponding to 75% VO2max and 80% maximal heart rate remained below the intensity at VT2, as was the average work rate during constant high intensity exercise (Table 2).

### Parameters of physical exercise during Exercising conditions

Each participant performed two Exercising conditions (apart from the rest session) at constant intensity in counterbalanced order. Gas exchanges were not assessed during exercising sessions. We tailored the intensities from the values obtained during the maximal incrementing effort test. We defined moderate intensity as 70% of maximal heart rate (corresponding to 60% of VO2max) and high intensity exercise as 80% of maximal heart rate (corresponding to 75% of VO2max). In the moderate intensity condition, participants exercised for 30 min at a work rate that was adjusted every three minutes to maintain heart rate around the individual target values corresponding to 70% of maximal heart rate. For the high intensity exercising condition, participants first warmed up for 3 minutes at a work rate corresponding to 60% of their maximal heart rate and then exercised for 15 min at a work rate that was adjusted to maintain HR at the individual target values corresponding to 80% of maximal heart rate. All predetermined and actual work rates are listed in Table 2.

For each participant, the work rates corresponding to 60% and 75% of VO2max and those corresponding to 70% and 80% of maximal heart rate were estimated, respectively on the individual VO2 / work rate and heart rate / work rate relationships, both obtained during the maximal incremental effort test for each participant. However, some discrepancy can occur between work rate at 60% VO2max and 70% maximal heart rate on one hand, and between work rate at 75% VO2max and 80% of maximal heart rate on the other hand. Consequently, we set the initial work rates, from exercise onset to the 3rd minute of constant exercise, based on the values estimated from the VO2 / work rate ratio. Then, after 3 minutes of exercise, and throughout the constant test, we adjusted the work rate to maintain the heart rate of the participant at the individually target values, respectively 70% and 80% of maximal heart rate, for moderate and intense exercise. We set up this regular adjustment to take into account the absence of steady-state of the VO2max protocol – which is inherent of incremental protocols – but from which submaximal work rates were estimated based on VO2 / work rate and heart rate / work rate relationships. We corrected for small heart rate drifts occurring during prolonged constant intensity exercises. But please note that, as reported in Table 2, the average work rates reported during both moderate and intense exercises remained very close to the initial predetermined work rate.

### Functional MRI data acquisition and analysis

A 3 Tesla MRI scanner (SIEMENS Trio System, Siemens, Erlangen, Germany) with a 32-channel head coil was used to acquire MRI data. We acquired T2*-weighted fMRI 2D images with a multiband echo-planar sequence, which acquires 3 slices at a time using axial slice orientation (66 slices; voxel size, 2 × 2 × 2 mm; repetition time (TR) = 1880 ms; echo time (TE) = 34 ms; flip angle (FA) = 60°). The last sequence of Visit 1 was a T1-weighted 3D sequence (192 contiguous sagittal slices; voxel size, 1.0 × 1.0 × 1.0 mm; TR = 1900 ms; TE = 2.27 ms; FA = 9°), which provided a whole-brain structural image.

We used SPM12 (Wellcome Department of Imaging Neuroscience, London, UK) for the analysis of the functional images. Preprocessing followed standard procedures: realignment, slice timing to correct for differences in slice acquisition time, normalization (to an MNI template), and smoothing (with an isotropic 8-mm FWHM Gaussian kernel). Corrections to regress out potential artifacts coming from heart rate and breathing were performed using Retroicor ^54^ and RVHcorr ^55, 56^.

We included reaction time (RT) as a parametric modulator of each trial. Specifically, we used the reaction time for each trial multiplied by −1 (or -RT), which allowed us to test for local increases in MRI signal linked to better performance (i.e. faster reaction times). Because moderate and high intensity exercise conditions did not differ in terms of their influence on performance or AEA levels, we considered both exercise conditions together and compared them to the rest condition. Thus, to reveal any effect of exercise on the parametric modulator modeling performance, we performed a contrast between exercise and rest (i.e. moderate and high exercise minus 2*rest), including all sequence blocks from the second session of every visit, and found activations in the right hippocampus and right caudate. Because we had strong a priori concerning the implication of the hippocampus and striatum in this task (see introduction section), we created corresponding regions of interest with the AAL atlas in the WFU PickAtlas toolbox version 2.4 ^57^, which we then used to perform small volume correction on the activation map.

As a quality check procedure, we tested whether sequence and random blocks elicited similar patterns of activity, as would be expected from the engagement of motor processes in both types of blocks. We thus ran a flexible factorial design on the contrast images corresponding to the main effects of each Block Type, using Subjects and Block Type (sequence, random) as factors. We then performed a conjunction analysis and obtained the common activations shown in Table 1 including right motor cortex and left cerebellum.

### Blood samples

Before and after each exercising condition, 2.5 mL of blood were collected into a BD Vacutainer K_2_EDTA 5.4 mg tube. This tube was immediately centrifuged for 10 minutes at 8009g at 4°C, the supernatant (plasma) was taken in aliquots of 200 μL. All samples were then frozen and stored at −80°C until analysis.

The endocannabinoid AEA levels were determined from 100 μl of plasma by liquid-liquid extraction. This was followed by liquid chromatography (Ultimate 3000RS, Dionex, CA, USA) and mass spectrometry using a 5500 QTrap triple quadrupole/linear ion trap (QqQLIT) mass spectrometer equipped with a TurboIon-Spray interface (AB Sciex, Concord, ON, Canada) as described previously ^58, 59^.

## ACKNOWLEDGEMENTS

We are grateful to the Brain and Behavior Laboratory of the University of Geneva (Geneva, Switzerland) for providing help and scanning facilities. We are particularly grateful to Dr. Tomas Ros for providing the scripts to successfully run and analyze the task.

## AUTHOR CONTRIBUTIONS

B.M.B., A.B., G.F., S.S., and K.I. designed research; B.M.B., A.B., M.G.L., N.I., and K.I performed research; B.M.B., A.B., M.G.L., E.L., A.T., S.S., and K.I. analyzed data; and all the authors wrote the paper.

## ADDITIONAL INFORMATION

The authors declare no competing interests. This work was supported by the Swiss National Science Foundation (No 320030_135653 to S. Schwartz, 32003B_127620 and 3200B0-114033 to G. Ferretti), the National Center of Competence in Research (NCCR) Affective Sciences financed by the Swiss National Science Foundation (No 51NF40-104897 to S. Schwartz). No conflicts of interest, financial or otherwise, are declared by the authors

